# CSF plasma cell expansion in LGI1-/CASPR2-autoimmune encephalitis is associated with loss of regulatory MAIT cells

**DOI:** 10.1101/2023.12.21.572754

**Authors:** Daniela Esser, Louisa Müller-Miny, Michael Heming, Manuela Paunovic, Martijn van Duijn, Ligia Abrante Cabrera, Katharina Mair, Christine Strippel, Saskia Räuber, Eric Bindels, Justina Dargvainiene, Heinz Wiendl, Sven G. Meuth, Jan Bauer, Nico Melzer, Maarten J. Titulaer, Frank Leypoldt, Gerd Meyer zu Hörste, EMC-AIE Study group

## Abstract

Anti-Leucine-rich glioma inactivated-1 (LGI1) and anti-contactin-associated-protein-2 (CASPR2) associated autoimmune encephalitis (AIE) variants are characterized by directly pathogenic autoantibodies present in serum and CSF. The dynamics and drivers of intrathecal and systemic autoantibody production are incompletely understood. We aimed to elucidate the immunologic basis of the LGI1-/CASPR2-associated AIE variants by performing multi-omic profiling of CSF/blood in untreated patients. We validated findings by flow cytometry in independent cohorts and confirmed functionality using rodent immunization.

We identified clonal IgG2 and IgG4 plasma cell expansion and affinity maturation in the CSF together with clonally restricted, activated, antigen-experienced CD8 and CD4 T cells as a hallmark of these encephalitis variants. Using recombinant cloning, we confirmed that expanded CSF plasma cell clones almost exclusively bound the respective neuronal autoantigen. In addition, we found a loss of regulatory mucosa-associated invariant T (MAIT) cells and gamma delta T cells in the CSF and – to a lesser degree – in blood. We validated the functional role of these invariant T cells using a novel murine active immunization paradigm using both autoantigens: MAIT cells suppressed systemic formation of LGI1 and CASPR2-specific anti-neuronal antibodies.

We propose that loss of systemic and intrathecal regulatory mechanisms mediated by innate-like T cells promote plasma cell expansion and autoantibody production as a shared mechanism in AIE.

**One sentence summary:** Cerebrospinal fluid (CSF) and peripheral blood (PB) single cell transcriptomics of patients with untreated anti-LGI1 and anti-CASPR2 autoimmune encephalitis demonstrated CSF specific expansion of autoantigen-specific plasma cell clones and systemic loss of invariant mucosa-associated T-cells (MAIT).

## Introduction

Anti-leucine-rich glioma inactivated 1 (LGI1) encephalitis and contactin-associated protein 2 (CASPR2) autoimmunity are two autoantibody-defined subgroups of rare, acquired inflammatory diseases affecting the central nervous system (CNS) (*1*). Together, they represent almost one fourth of all antibody-positive autoimmune encephalitis (AIE) patients (*2–4*). They are clinically and epidemiologically distinct: LGI1-AIE affects older individuals of both genders while CASPR2-autoimmunity mainly occurs in elderly men. LGI1-AIE often manifests with acute onset focal seizures - including facio-brachio-dystonic seizures - eventually evolving into limbic encephalitis (memory dysfunction, psychiatric symptoms and epileptic seizures) (*1, 5*). Anti-CASPR2 autoimmunity has a more heterogeneous manifestation and can affect the central, peripheral and autonomic nervous system with limbic encephalitis, ataxia, movement disorders, sleep abnormalities, neuropathic specific pain, peripheral nerve hyperexcitability syndromes, autonomic dysfunction and weight loss (*6*). Of note, almost all patients have core features of limbic encephalitis (90%) and cause considerable disability.

Initiation and propagation of the underlying autoimmune reaction are incompletely understood. However, both diseases share immunological features suggesting related immunological mechanisms: (1) A strong human leukocyte antigen (HLA) class II association (*2, 6, 7*) indicating T-cell involvement. (2) Autoantibodies mostly of the IgG4 isotype (*8*). These non-complement-binding non-crosslinking IgG4 autoantibodies disrupt their target antigens by steric interference with protein function (*8, 9*). However, it remains unclear what drives the IgG4 isotype switch in these diseases, a process also requiring T cell help. Notably, LGI1- and CASPR2-AIE lack overt inflammatory changes in the CSF. The median leukocyte concentration in CSF in LGI1-AIE is 1.5/µl, only 11% of patients have mild lymphocytic pleocytosis, and <20% of patients show CSF restricted oligoclonal bands (*3*) with similar results in CASPR2 autoimmunity (*10, 11*). This peculiar absence of inflammatory findings in routine CSF studies has been interpreted as an argument against a relevant intrathecal component of the misled adaptive autoimmune reaction. However, in light of the often predominantly systemic effect of many immunomodulatory strategies, e.g. plasma exchange and depletion of CD20-B-cells, there is a mechanistic, diagnostic and therapeutic need to clarify the role of intrathecal, compartmentalized antigen-specific immune reactions in these diseases.

Here, we aimed at better understanding the role of the intrathecal immune compartment in comparison to the systemic immune reaction in two model diseases defined by adaptive autoimmunity against defined neuronal autoantigens: LGI1- and CASPR2-AIE. We performed a cross-compartment immune cell characterization of large prospectively recruited, multi-center cohorts of untreated LGI1-/CASPR2-AIE-patients by integrating single cell RNA-sequencing and flow cytometry of CSF and blood. We identified a strong intrathecal expansion of IgG2 and IgG4 class-switched B lineage plasmablasts and plasma cells with signs of intrathecal affinity maturation together with clonally expanded, activated CD4- and CD8-T-cells as a shared feature of LGI1-AIE and CASPR2-AIE. Almost all recombinant antibody clones generated from highly expanded plasma cells from the CSF of LGI1-AIE and CASPR2-AIE patients consistently recognized their corresponding neuronal target antigen and showed strong signs of somatic hypermutation. In addition, we found and confirmed a loss of immune-regulatory mucosa-associated invariant T (MAIT) cells in CSF and in the peripheral blood of LGI1-/CASPR2-AIE that suppressed specific autoantibody formation in a rodent active immunization model. We thus identified autoantigen-specific intrathecal B cell expansion and ongoing affinity maturation as well as peripheral loss of immune-regulatory mechanisms as converging mechanisms in two common variants of autoimmune encephalitis: LGI1- and CASPR2-AIE.

## Results

### Sample acquisition, patient and CSF characteristics of LGI1-AIE and CASPR2-AIE patients

We prospectively recruited, treatment-naive patients with LGI1-AIE (CSF: n=8; blood n=7) and CASPR2-AIE (CSF: n=5, blood n=3) across three clinical centers (cohort #1). We directly *ex vivo* analyzed CSF and blood cells acquired in parallel using single cell RNA-sequencing (scRNA-seq) (Fig. 1A) as established previously (*12*). Sample processing was standardized between centers to minimize systematic technical bias (see methods). Clinical features of patients were representative of the known phenotype of LGI1-/CASPR2-AIE (Suppl. Tab. 1). Two of eight LGI1-AIE and zero of five CASPR2-AIE patients were female; the average age of the combined group was 63 (95% CI: 60-67) (Suppl. Tab. 1) (*1*). Basic CSF analyses (e.g. cell numbers, protein) did not show gross inflammatory changes and basic CSF parameters also did not significantly differ between LGI1- and CASPR2-AIE (Suppl. Fig. 1A) in accordance with previous studies (*3, 10*). As controls, we included prospectively recruited (n = 5) patients with idiopathic intracranial hypertension (IIH) (Suppl. Tab. 1) and existing published data from one center (n = 9; (*12, 13*)).

**Fig. 1.**
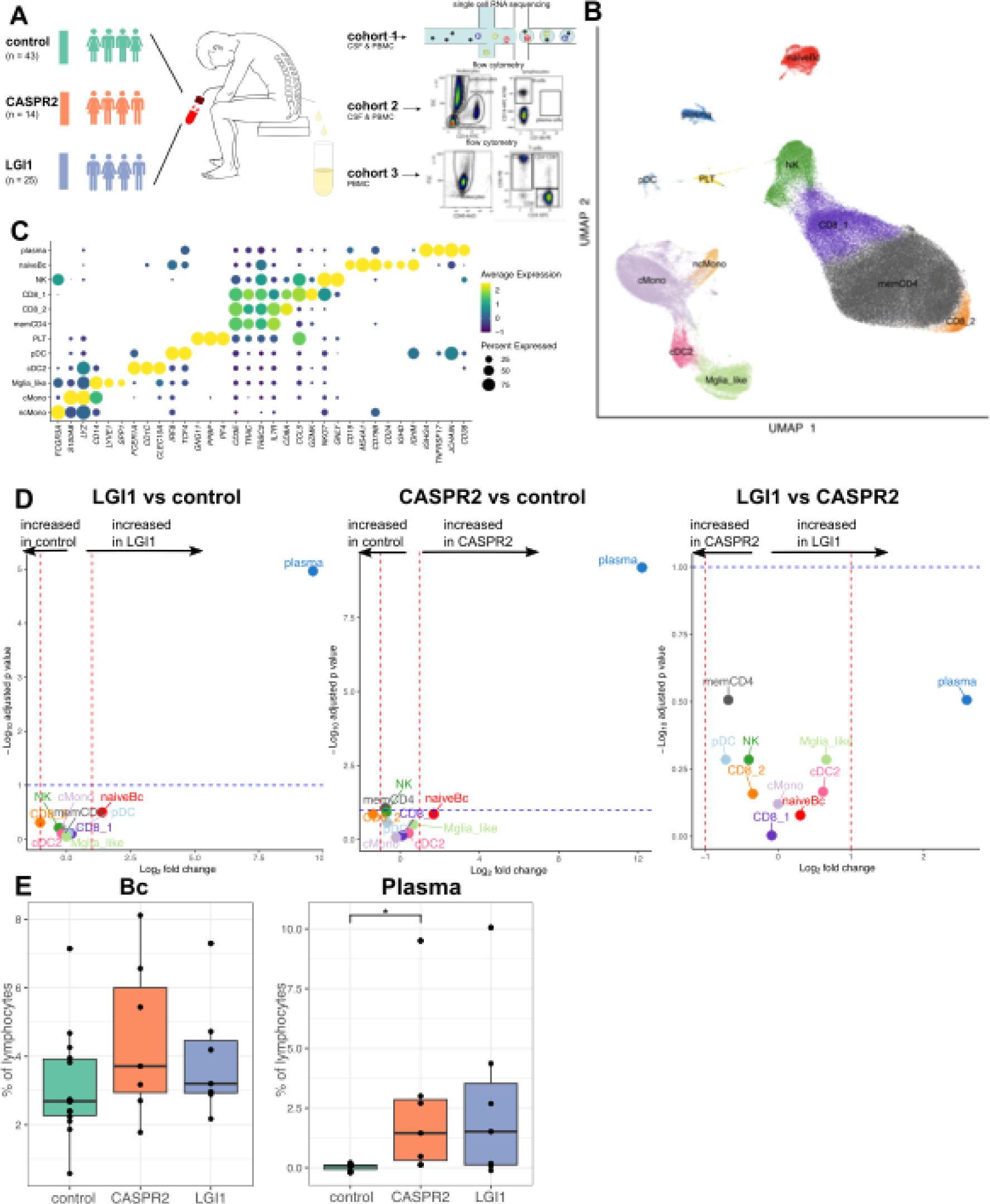
Single cell transcriptomics and flow cytometry identifies expansion of plasma cells as hallmark of LGI1-/CASPR2-AIE. (A) Schematic representation of sample cohorts, data processing and analysis. Graphic adapted from (*13, 86*) (B) Uniform Manifold Approximation and Projection (UMAP) plot showing 12 color-coded cell clusters of 68,459 single cell transcriptomes integrated from cerebrospinal fluid (CSF) and PBMC cells from LGI (n = 8) and CASPR2 (n = 5) patients, and idiopathic intracranial hypertension controls (control = 14). (C) Dot plot depicts selected marker genes. Color encodes mean expression, dot size visualizes the fraction of cells expressing the gene. (D) Volcano plots depicting differentially abundant clusters in LGI1 vs. control (left panel), CASPR2 vs. control (middle panel), and LGI1 vs. CASPR2 (right panel). The log2 fold change of the differential abundance is plotted against the negative log10 of the adjusted p values. The threshold for the p value was set at 0.1 and for the fold change at 2. (E) CSF cells of cohort 2 were analyzed by flow cytometry. Relative percentage of B cells (Bc; %CD3^-^CD19^+^CD138^-^) and of plasma cells (Plasma; %CD3^-^ CD19^+^CD138^+^) were quantified as percentages of all lymphocytes and visualized in box plots. (Suppl. Fig. 2A). Boxes represent lower quartile, median and upper quartile. Whiskers include 1.5 times the interquartile range. Statistical significance determined by Kruskal-Wallis with post hoc Dunn’s test and Benjamini-Hochberg adjusted. Cluster names: plasma: plasma cells; naïveBc: naïve B cells; CD8: CD8+ T cells; memCD4: memory CD4+ T cells; PLT: platelets; pDC: plasmacytoid dendritic cells; cDC: conventional dendritic cells; Mglia_like: microglia like cells; NK: natural killer cells; cmono: classical monocytes; ncmono: non classical monocytes

### CSF plasma cell expansion is a hallmark of LGI1-AIE and CASPR2-AIE

We next analyzed CSF cells and peripheral blood mononuclear cells (PBMC) by scRNA-seq in an unbiased fashion. We merged all scRNA-seq data from CSF and PBMC of AIE and control patients, which encompassed 162,360 total single-cell transcriptomes (CSF: 75,496; blood: 86,864) with 3,690 mean ± 505 SEM cells per sample and 1,026 mean ± 118 SEM genes detected per cell (Suppl. Tab. 2). Single cell transcriptomes are henceforth termed ‘cells’ for simplicity. We performed cell clustering (Fig. 1B) and semi-automatically annotated the resulting 12 cell clusters (Fig. 1C; Suppl. Tab. 3). Across patients and controls, T cells (CD4^+^ > CD8^+^) and myeloid lineage cells dominated in CSF as described previously (*12*) (Suppl. Fig. 1B-D). However, when comparing LGI1- and CASPR2 encephalitis patients to controls, we found a pronounced increase of the CSF plasma cell (expressing *CD19, CD38, IGHG4*) and B cell clusters both in LGI1-AIE and in CASPR2-AIE patients compared to controls (LGI1>CASPR2, Fig. 1D). Proportions of other CSF cell lineages were not significantly altered compared to controls and in between encephalitis variants (Fig. 1D; Suppl. Tab. 4). Analysis of blood cells also did not show significant cell type differences in any comparison (data not shown). In summary, expansion of plasma cells represented the specific compositional hallmark of CSF in LGI1-/CASPR2-AIE.

### Flow cytometry confirms CSF plasma cell expansion in patients with LGI1-/CASPR2-AIE

Next, we aimed to confirm these transcriptomics-based observations. We were unable to prospectively perform parallel CSF scRNAseq and flow cytometry experiments due to severely limited available cell numbers in CSF for each technique. Therefore, we retrospectively identified independent cohorts of treatment-naive LGI1-AIE (n = 7) and CASPR2-AIE (n = 7) patients (cohort #2) whose CSF cells had been analyzed by multicolor flow cytometry as part of the routine clinical work-up in one center (Methods) (Suppl. Tab. 1). Patients with somatoform disorders (n = 14) served as non-inflammatory controls and were matched in age and gender (Suppl. Tab. 1). Basic clinical and CSF parameters of cohort #2 were similar to cohort #1 and replicated the known features of LGI1-/CASPR2-encephalitis patients (Supp. Tab. 1, Suppl. Fig. 2A-C). In this retrospective CSF flow cytometry data set, we confirmed the strong expansion of plasma cells (CD3^-^CD19^+^CD138^+^) in the CSF of LGI1-/CASPR2-AIE (Fig. 1E).

### CSF plasmablasts and cells are clonally expanded and express IgG2 and IgG4 isotypes

Accordingly, we next sought to characterize the CSF B lineage cells and specifically plasma cells in LGI1- and CASPR2-AIE. We sub-clustered all single cell transcriptomes annotated as B lineage from the merged PBMC/CSF scRNA-seq dataset. These B lineage cells separated into plasma cell/blast clusters (antibody-secreting cells, ASCs) and a naive/memory B cell cluster (Fig. 2A-C) as supported by their respective expression profile (Suppl. Fig. 3A). Notably, the ASC cluster dominantly contained CSF-derived cells from patients with AIE (Fig. 2A), with no differences between AIE variants and a high frequency of clonally expanded cells (Fig. 2B). Annotating ASC using a semi-supervised approach (Azimuth; Methods) and a tonsil reference (*14*), we successfully categorized most CSF cells within these clusters as either IgG plasma cell precursors, plasmablasts or less frequently mature IgG plasma cells. These were significantly increased in AIE-derived CSF cells compared to PBMC (Fig. 2C, D, Suppl. Fig. 3B), with distinct expression profiles (Suppl. Fig. 3C, D). B cell receptor sequence analysis identified that ASC transcribed IgG4 heavy chain genes in CSF cells significantly more often than in blood-derived cells and showed a tendency towards higher IgG2 abundance in CSF (Fig. 2E, F). Comparing transcribed isotypes in blood-derived plasma cells/memory-B cells, we did not observe a significant difference between patients (Suppl. Fig. 3E) and controls (not shown). Between LGI1- and CASPR2-AIE, cluster abundance and isotype gene expression did not differ, Suppl. Fig.3 B,E). We also did not observe an expansion of non-class switched cells or IgA or IgM producing cells in CSF (Fig. 2E). In summary, LGI1- and CASPR2 encephalitis patients harbor clonally expanded plasma cells and plasmablasts in their CSF, that preferentially transcribe IgG2 and IgG4 heavy chain genes.

**Fig. 2.**
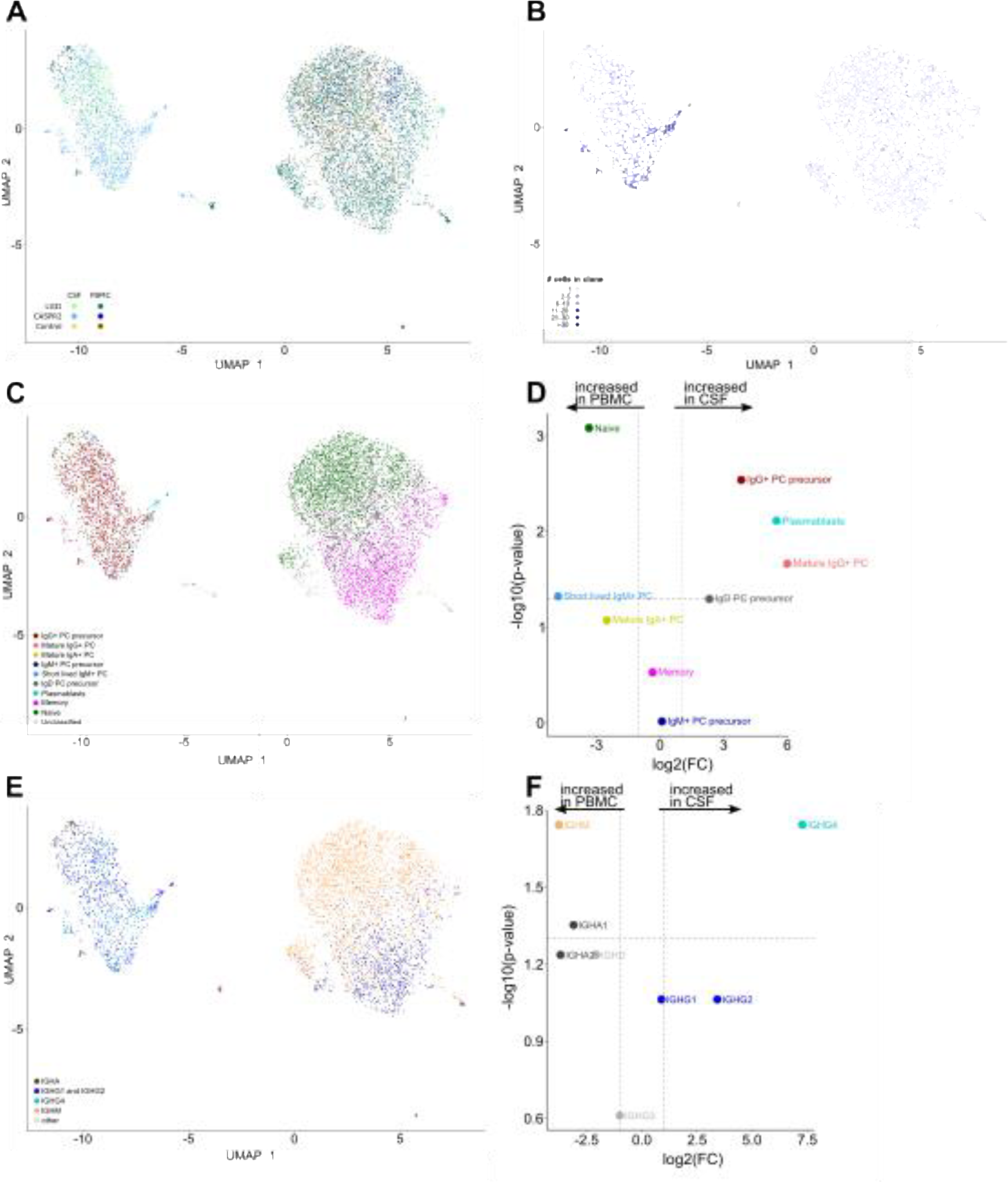
Plasmablasts and plasmacells are preferentially CSF-derived, clonally expanded, and express IgG2 and IgG4 heavy chains in LGI1-/CASPR2-AIE. Single cell transcriptomes in CSF (n = 2352, light colors) and PBMC (n = 4987, dark colors) identified as B cells from LGI1 (green, n = 8), CASPR2 patients (blue, n = 5), and idiopathic intracranial hypertension controls (IIH, brown, n = 14) were sub-clustered. Uniform Manifold Approximation and Projection (UMAP) plot with B lineage subclusters plasma blasts, plasma cells (PC) and memory and naive B cell indicated. Selected marker genes shown in Suppl. Fig. 3A. (A) UMAPs highlighting the sample type (PBMC vs. CSF) and cohort. (B) UMAP illustrating the distribution of clonally expanded B cells. All cells with B cell receptors occuring more than once were defined as clonally expanded (clone size indicated by shade). Clonotypes were based on heavy and light chain VDJ nucleotide sequence similarity and CDR3 length (algorithms by Cell Ranger, see methods). (C) UMAP depicting Azimuth annotations using a tonsil reference to differentiate plasma blasts from plasma cell precursors and mature plasma cells and by isotype. (D) Volcano plot depicting differential relative proportion differences between PBMC (left) and CSF (right). The log2 fold change (FC) of the differential abundance is plotted against the negative log10 of the adjusted p values.(E) UMAP depicting the preferentially expressed immunoglobulin heavy chain genes. (F) Proportions of B cells dominantly expressing the respective IGH gene were calculated for each patient and differences between PBMCs (left) and CSF (right) illustrated in the volcano plot. Only samples with at least 50 cells were considered.

### Clonally expanded CSF plasma cells almost exclusively recognize the disease-defining autoantigen in LGI1- and CASPR2-patients

Next, we aimed to differentiate LGI1-/CASPR2-autoantigen-specific plasma cells in CSF from potential non-antigen-specific “bystanders”. For a representative subgroup of patients (n=8) from cohort #1, we bioinformatically identified the top 5-9 largest expanded B lineage clones and identified those with concurrently available consensus heavy and light chain BCR variable region sequence information (LGI1: n = 27 from 5 patients, CASPR2: n = 15 from 3 patients). We synthesized and cloned the corresponding variable heavy and light chain sequences into human IgG4 constant heavy and kappa or lambda light chain vectors, produced and affinity-purified the recombinant human monoclonal antibodies (rhumAbs). Out of 27 rhumAbs, 25 (93%) bound their respective target antigen on the surface of transfected HEK293T-cell-lines thus confirming the autoantigen-specificity of the vast majority of expanded clones across patients and AIE variants ((*15–17*), Suppl Fig. 4A). We next visualized the CSF and blood B cell repertoire using clone trees (including but not limited to the experimentally validated clones) for all patients. Exemplary graphs are shown for one LGI1 and one CASPR2 encephalitis patient (Fig. 3A-B). As described for CSF, B cell lineage cells were almost entirely absent from control CSF and could not be used as controls, therefore findings were compared between CSF and blood B cell compartments. In general, we observed a highly restricted, clonally expanded B cell repertoire in the CSF of CASPR2 and LGI1 patients while mostly singletons and only rare single expanded clones were detected in blood. There were almost no shared clones between CSF and blood (Fig. 3A-B). CSF clones showed a high degree of clonal expansion and somatic hypermutations which was more pronounced in the CASPR2 cohort (Fig. 3C-E, Suppl. Fig. 4B) and showed reduced diversity compared to blood B cells (Suppl. Fig. 4C-D). Of note, even though most expanded clones consisted of plasma cells and plasma blasts, some CSF clones also contained class-switched memory-B cells (Fig. 3F). B cell clones were mostly expanded with identical CDR3 sequences, however, many CSF B cell clones also showed within-clone mutations (Suppl. Fig. 4E-F), predominantly in CASPR2 AIE patients.

**Fig. 3.**
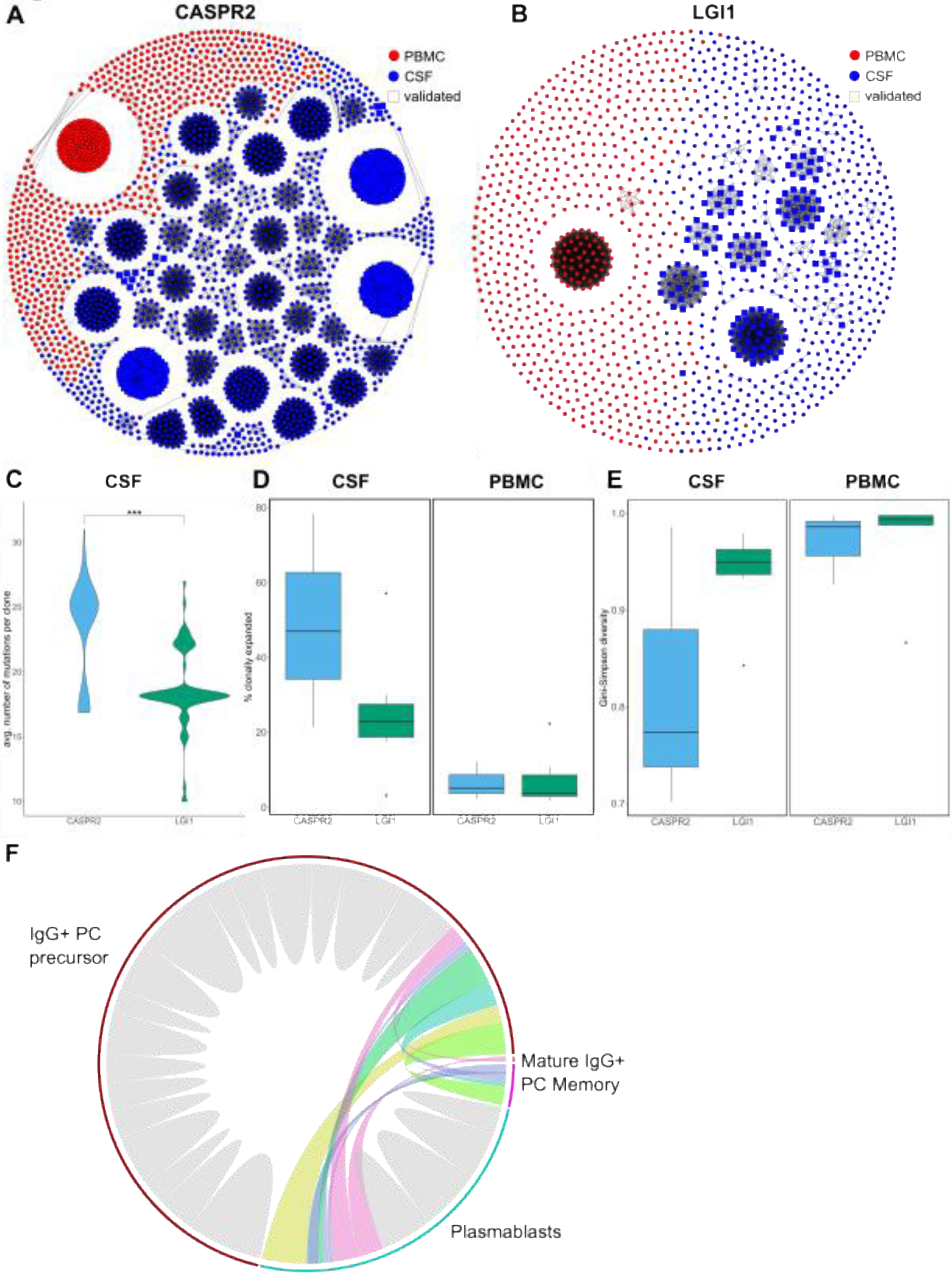
Clonal restriction and expansion with ongoing affinity maturation within B cell clones in the CSF of LGI1-/CASPR2-AIE. (A, B) Visual representation of CSF (blue) and blood (red) B cell receptor networks in two representative patients with LGI1-AIE (A) and CASPR2-AIE (B). Each dot indicates one cell. Lines connect cells if they share the same genes in the variable, joining, and constant segments and have identical CDR3 regions defined as zero mismatches in the nucleotide sequence. Squared symbols indicate clone sequences experimentally verified as autoantigen-specific using recombinant human antibodies (Suppl. Fig. 4D). (C) Percentage of clonally expanded cells, as defined as CDR3 sequence present in >1 cell. (D) Gini-Simpson index as a measure of clonal diversity compared between AIE types and compartments. (E) Mean number of mutations in the nucleotide sequences of each productive B cell clone determined as differences to the most similar IMGT reference sequence. (F) Circos plot highlighting clones shared between cells annotated as different B cell types. Clones within one cell type are marked in gray, clones shared between cell types are shown by colored connecting lines. Line thickness indicates the number of cells; B cell clusters named as in Figure 2C.

This demonstrates that a restricted B cell repertoire in CSF contains highly expanded ASC in patients with LGI1-AIE and CASPR2-AIE, which consistently recognize their corresponding neuronal target antigen. Autoantigen-specific memory B cells also exist in CSF and ongoing extra-germinal center affinity maturation occurs locally within the CSF.

### Clonally expanded CD4 and CD8 T cells are activated and partially exhibit B cell-helper capacity in the CSF

We next characterized T cells in the CSF of LGI1- and CASPR2-AIE and their potential contribution to local B cell expansion. We first sub-clustered and annotated all single cell transcriptomes identified as T cells (see methods) (Fig. 4A). As described (*12*), T cells spanned a transcriptional gradient ranging from naive towards memory phenotype (Fig. 4A,B) rather than forming distinct subclusters. Next, we analyzed transcriptional differences between AIE and control patients. The number of differentially expressed (DE) genes in the AIE vs. control comparison was highest in memory CD8 and CD4 T cells with transcriptional signs of activation (Suppl. Tab. 5,6). This was also reflected in a cluster-free approach for localizing transcriptional alterations (*18*) that preferentially localized DE genes into neighborhoods of memory T cells (Fig. 4C). CSF-localized CD4^+^ and CD8^+^ T cells are thus antigen-experienced and activated in LGI1-/CASPR2-AIE patients.

**Fig. 4.**
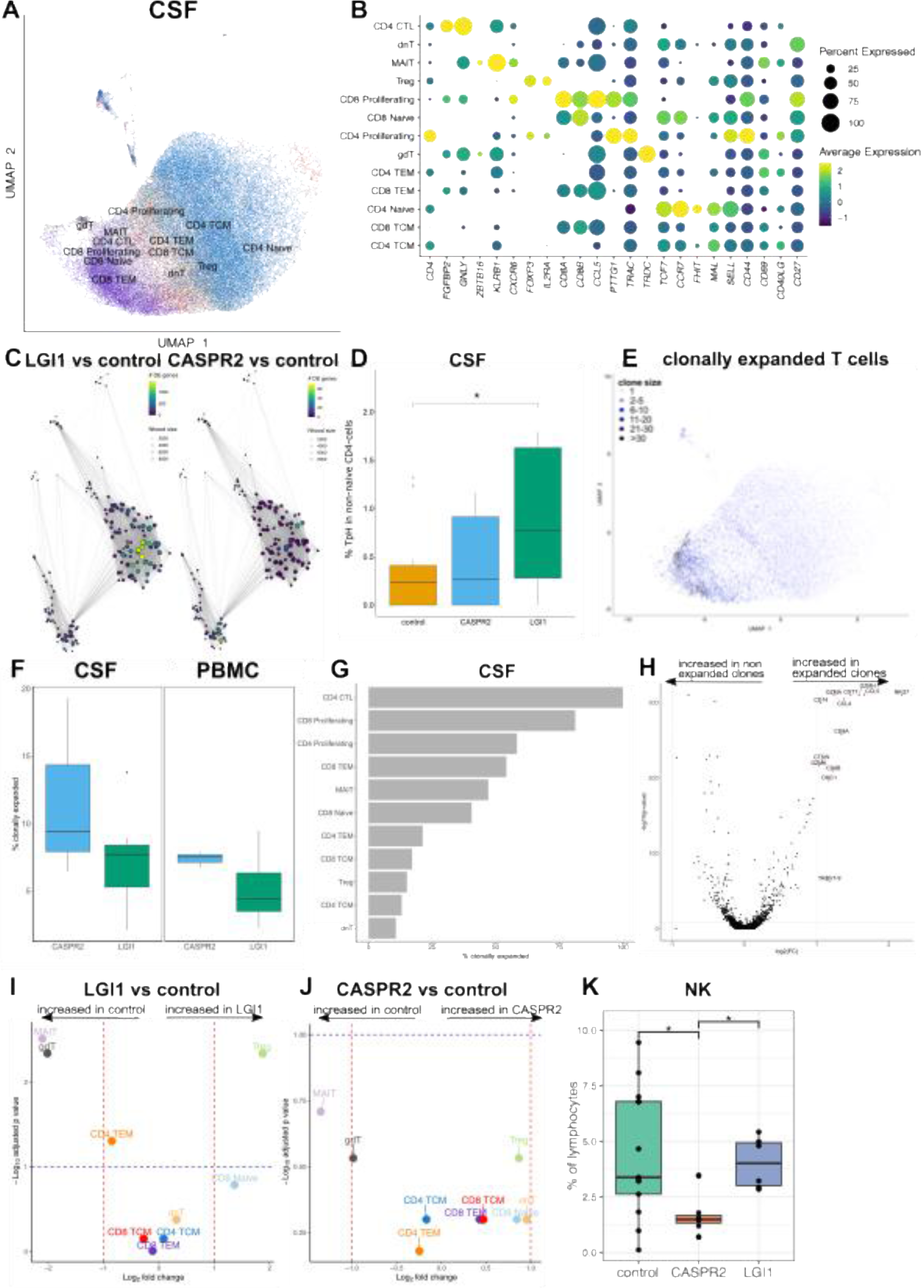
Clonal expansion of activated CD4 and CD8 T cells and loss of innate-like MAIT cells in the CSF of LGI1-/CASPR2-AIE. (A) CSF cells identified as T cells, were sub-clustered from LGI (n = 8) and CASPR2 patients (n = 5), and idiopathic intracranial hypertension controls (control = 14). Uniform Manifold Approximation and Projection (UMAP) plot showing 13 color-coded T cell subclusters. (B) Dot plot showing selected marker genes of the clusters. The color encodes the mean expression, dot size visualizes the fraction of cells expressing the respective gene. (C) Transcriptional neighborhood analysis (miloDE) depicting differentially expressed genes in LGI1 vs. control (left panel) and CASPR2 vs. control (right panel) in all 13 parent clusters. The color encodes the number of differentially expressed (DE) genes, dot size visualizes neighborhood size. (D) Boxplot depicting percentage of TpH cells (*PDCD1*/PD-1^+^, *PRDM1*/BLIMP-1^+^, CXCR5^-^) of non naive CD4-cells in the CSF across all three cohorts (E) T cell subcluster UMAP visualizing clone frequency in the CSF of LGI1-/CASPR2-AIE patients. (F) Boxplot showing the percentage of clonally expanded T cells for CASPR2- and LGI1-AIE split by sample material (G) Percentage of clonally expanded CSF cells from AIE patients separated by cell type. (H) Volcano plot showing differential expression between expanded and non expanded T cell clones. The log2 fold change of the differential abundance was plotted against the negative log10 of the adjusted p values. (I) Volcano plots depicting differential abundance in the CSF of T cell sub-clusters in LGI1 vs. controls (I) and CASPR2 vs. controls (J). The log2 fold change of the differential abundance was plotted against the negative log10 of the adjusted p values. (K) In an independent flow cytometry cohort (#2), NK cells (CD3^-^CD56^+^) were reduced in the CSF of CASPR2-AIE patients. Boxplot depicting percentage of all lymphocytes in patients with AIE and controls. Abbreviations/cluster names: CD4 CTL, dnT: double negative T cells, MAIT: mucosal-associated invariant T cells; Treg: regulatory T cells, CD8+ Proliferating: proliferating CD8 T cells, CD8 Naive: naive CD8+ T cells, CD4 Proliferating: proliferating CD4+ T cells; gdT: gamma delta T cells, CD4 TEM: CD4+ effector memory T cells; CD8 TEM: CD8+ effector memory T cells, CD4 Naive: naive CD4 cells, CD8 TCM: CD8+ central memory T cells, CD4 TCM: CD4+ central memory T cells, NK: natural killer cells, TpH: peripheral T helper cells

We next queried our data for T cells with B cell-helping capacity such as peripheral T helper (TpH) cells. Indeed, a small (∼1%) proportion of CD4^+^ T cells with a TpH-like transcriptional phenotype (*PDCD1*/PD-1^+^, *PRDM1*/BLIMP-1^+^, CXCR5^-^) was detected in the CSF. Their proportion in CSF was significantly increased in LGI1-AIE with a similar trend in CASPR2-AIE in comparison to controls (Fig. 4D). Interaction network analysis of CSF B/T cells confirmed memory CD4 T cells (CD4 TCM and CD4 TEM) to be central interacting cells within the lymphocyte compartment (Suppl. Fig. 4G). Gene set enrichment analysis (GSEA) based on all non-naive T cells in the CSF suggested a strong activation of the T cell receptor signaling pathway activation and antigen processing/presentation (Suppl. Fig. 4H). T cell receptor (TCR) repertoire analysis showed prominent clonal expansion and restriction in CSF, with multiple clones spanning CSF and blood compartments (Suppl. Fig 4I). Expanded clones were often localized in the region of proliferating CD4 (CD4CTL) and CD8 T cells regions (Fig. 4E-G). Differential expression analysis of expanded vs. non expanded CSF clones highlighted an upregulation of cytotoxic and activation T cell genes (*GZMK, GZMA, CCL4, CCL5*) (Fig. 4H). The CDR3 motifs of the TCR-β chain were partially shared among patients. One TCR motif (TGDSNQP) was observed in 4 out of 8 LGI1 CSF samples, but only in 2 out of 26 other CSF repertoires (p=0.01). The TCR motif was evaluated against VDJdb, a database of known TCR epitope associations, and revealed a connection to Epstein Barr Virus (*19*). Overall, activated clonally expanded T cells with partially shared CDR3 motifs may thus contribute to affinity maturation of autoantigen-specific plasma cells/blasts in LGI1-/CASPR2-AIE and with a potential link with preceding infections.

### Innate-like T cells with regulatory phenotype are lost from the CSF in LGI1- and CASPR2-AIE

Next, we aimed for an unbiased compositional analysis of CSF T cells. When systematically comparing the proportions of T cell sub-clusters, we surprisingly identified a reduction in the proportion of cells annotated as MAIT (e.g. *CXCR6*, *KLRB1*/CD161, *ZBTB16*; Fig 4B) and γδ T cells (cluster named gdT; e.g. *TRDC*) in CSF, which reached the significance threshold in LGI1-AIE, but not in CASPR2-AIE compared to controls (Fig. 4I,J); likely due to differences in statistical power (n=8 vs. n=5). Notably, MAIT cells sense bacterial products and have been ascribed regulatory function and loss of MAIT cells has been associated with multiple autoimmune diseases (*20*).

We therefore next sought to confirm the loss of innate-like T cells (including MAIT cells) in AIE in independent cohorts. We first queried the flow cytometry data of CSF retrospectively collected in cohort #2 for differences in such lineages. Although no comprehensive profiling of CSF and no specific markers of MAIT cells were retrospectively available, we found that the proportion of CD3^-^CD56^+^ NK cells in the CSF was significantly lower in CASPR2-AIE compared to control patients (Fig. 4K). Other cellular parameters assessed by flow cytometry did not differ between AIE subtypes and compared to controls (Suppl. Fig. 2C). Together with NK and γδ cells, MAIT cells belong to the category of innate-like T cells. We therefore speculated that loss of innate-like T cells could contribute to autoantibody formation in LGI1/CASPR2-AIE and possibly generally in humoral autoimmunity.

### Loss of innate-like T and MAIT cells in the peripheral blood in LGI1-/CASPR2-AIE

Next, we investigated whether the loss of innate-like T-cells of patients with LGI1- or CASPR2-AIE was restricted to the CSF compartment or also evident in the systemic immune compartment. We first analyzed scRNA-seq data from blood T cell sub-clustering (Fig. 5A). When comparing both AIE subtypes, we identified a significant reduction of clusters annotated as CD4 cytotoxic T cells in both LGI1- and CASPR2-AIE vs. controls and of gdTc in CASPR2-AIE vs controls but not LGI1-AIE as well as a tendency for a reduction of MAIT cells in CASPR2-AIE (Fig. 5B). The reduction of γδ T cells is thus maintained across CSF and blood compartments in CASPR2-AIE. The relative proportion of the MAIT cell cluster was not significantly different; potentially due to lack of power or low absolute proportions of MAIT cells. Therefore, we aimed to confirm this in an independent, larger cohort. We characterized T cells including γδ T cells and MAIT cells by flow cytometry in available cryo-preserved blood mononuclear cells from an additional cohort of LGI1-AIE (n = 14), CASPR2-AIE (n = 11) and matched controls (n = 14) (cohort #3) (Methods, Suppl. Fig. 5A). This cohort also featured clinical and CSF features in accordance with the disease (Suppl. Tab. 1). We confirmed a significant decrease of blood MAIT cells, but not of γδ T cells and other T cell subsets, in LGI1-and CASPR2-AIE compared to controls (Fig. 5C,D, Suppl. Fig. 5B). Cross-compartment loss of innate-like T-cells (including MAIT cells) is thus a shared feature of both LGI1-/CASPR2-AIE subtypes.

**Fig. 5.**
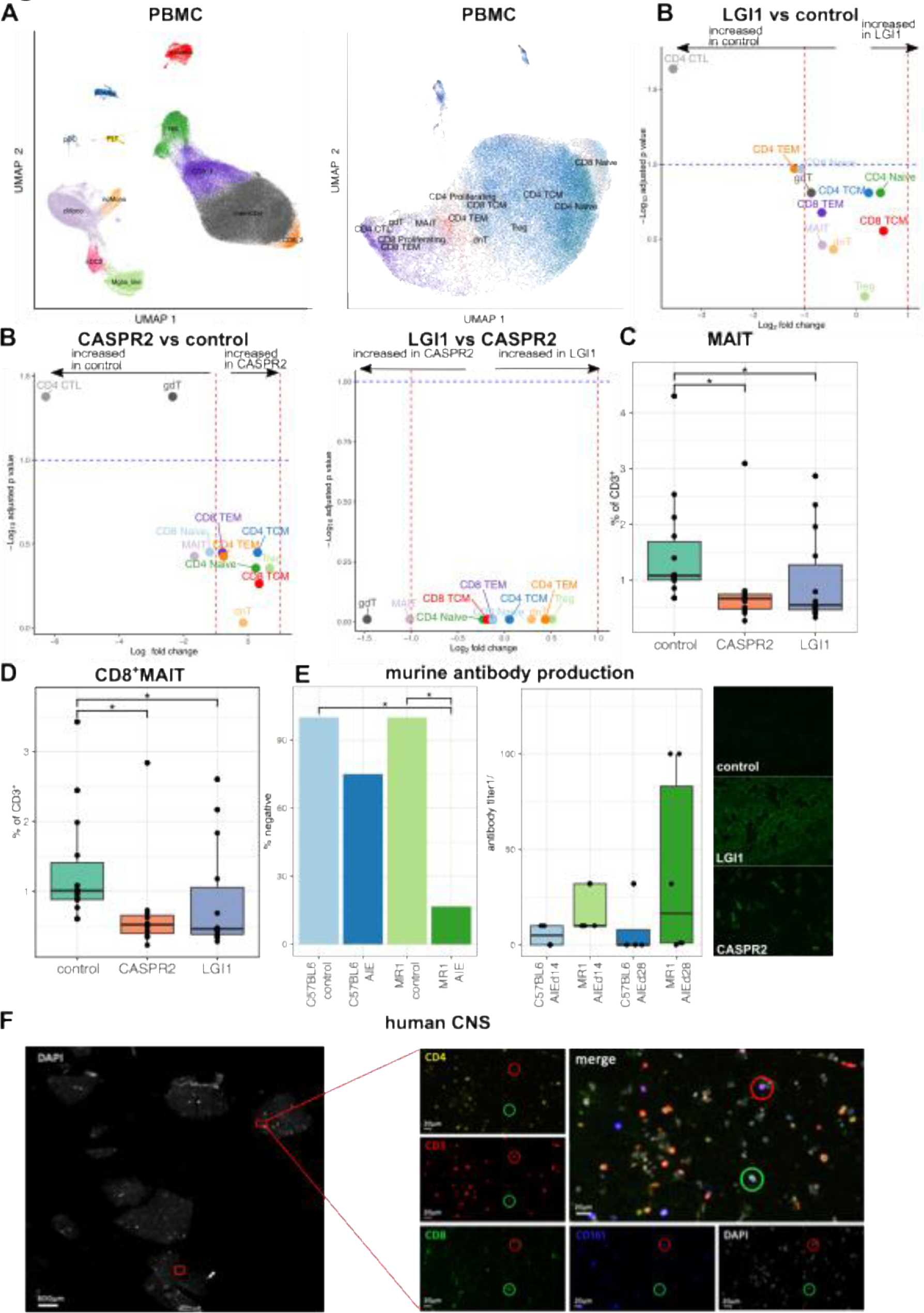
Peripheral MAIT cells are lost in LGI1-/CASPR2-AIE and suppress autoantibody generation in a murine active immunization model. (A) Uniform Manifold Approximation and Projection (UMAP) plot shows cell clusters of all PBMCs (left panel) and cell types of all PBMC T cells (right panel) from LGI and CASPR2 patients (LGI1 n = 7, CASPR2 n = 3), and idiopathic intracranial hypertension controls (control = 14). (B) Volcano plots depict differentially abundant clusters in LGI1 vs. control (left panel), CASPR2 vs. control (middle panel) and LGI1 vs. CASPR2 (right panel) of T cell subclusters. The log2 fold change of the differential abundance is plotted against the negative log10 of the adjusted p values. (C+D) PBMCs analyzed by flow cytometry of cohort #3. The proportion of (C) MAIT cells and (D) CD8+ MAIT cells among all T cells in patients with AIE is visualized in box plots and categorized by disease group. (Suppl. Fig. 7B). The boxes represent the lower quartile, median and upper quartile. Whiskers include 1.5 times the interquartile range. Statistical significance was determined by Kruskal-Wallis with post hoc Dunn’s test and Benjamini-Hochberg adjusted.(E) Serum Antibody incidence (left) and antibody titer (middle) of mice serum on day 28 as well as representative images of indirect immune fluorescence (right, rodent sera on LGI1 and CASPR2 transfected cell lines). Statistical significance was determined by Fisher’s exact test. (F) Representative immunofluorescence of post-mortem histology (hippocampus, uncus) of a patient with anti-VGKC-AIE. Innate T cell-like cells were identified via CD3^+^CD4^+^CD8^+^CD161^+^ and CD3^+^CD4^+^CD8^-^gdTCR^+^ markers. Abbreviations: plasma: plasma cells; naïveBc: naïve B cells; CD8: CD8^+^ T cells; memCD4: memory CD4^+^ T cells; PLT: platelets; pDC: plasmacytoid dendritic cells; cDC: conventional dendritic cells; Mglia_like: microglia like cells; NK: natural killer cells; cmono: classical monocytes; ncmono: non classical monocytes; CD4 CTL, dnT: double negative T cells, MAIT: mucosal-associated invariant T cells; Treg: regulatory T cells, CD8+ Proliferating: proliferating CD8 T cells, CD8 Naive: naive CD8^+^ T cells, CD4 Proliferating: proliferating CD4^+^ T cells; gdT: gamma delta T cells, CD4 TEM: CD4^+^ effector memory T cells; CD8 TEM: CD8^+^ effector memory T cells, CD4 Naive: naive CD4 cells, CD8 TCM: CD8^+^ central memory T cells, CD4 TCM: CD4^+^ central memory T cells

### MAIT cells infiltrate the brain in AIE and suppress peripheral anti-neuronal humoral autoimmunity

We next aimed to functionally confirm that MAIT cells repress systemic humoral anti-LGI1-/CASPR2-autoimmunity. However, available animal models of LGI1-/CASPR2-AIE are induced by passive xeno-transfer of autoantibodies and therefore only mimic effector mechanisms of antibodies (*21*). We therefore actively immunized mice with the recombinant extracellular portion of LGI1 and CASPR2 proteins (Methods; Suppl. Fig. 6A) adopting an immunization scheme previously successful in modeling anti-NMDAR AIE (*22*). After 15 days, 25% (1 of 4) of C57BL6 mice showed detectable LGI1 / CASPR2 autoantibodies in serum (Fig. 5E). We concurrently immunized littermates genetically deficient in the invariant MR1 antigen presenting molecule that consequently lack MAIT cells (*23*). Immunization (3 CASPR2, 3 LGI1) induced antineuronal serum antibodies in 83,3% (5 of 6) of MR1-deficient mice (Fig. 5E). Anti-LGI1-/CASPR2 antibodies remained undetectable in unimmunized mice from both genotypes (n = 10; not shown). In addition to antibody prevalence, antibody titers were also higher in MR1-deficient mice than in MR1-competent controls (Fig. 5E). Employing a novel active immunization approach, we thus demonstrate that loss of MAIT cells facilitates peripheral anti-neuronal humoral autoimmunity.

We next tested whether induction of peripheral anti-LGI1-/CASPR2-antibodies resulted in CNS infiltration and clinical signs of AIE in these mice. We performed flow cytometry of gross CNS-infiltrating leukocytes, but were unable to find numerical or compositional changes (Suppl. Fig. 6B). We next performed behavioral testing using a combination of paradigms to test for anxiety (3-chamber test, open field test) and learning and memory (novel object recognition). Again, we found no apparent phenotype and no significant differences in the behavioral test results between immunized and non-immunized mice of any genotype (Suppl. Fig. 6C). This indicates that peripheral anti-LGI1-/CASPR2-antibodies do not suffice to induce clinically and analytically tangible AIE-like disease in mice. This could be due to insufficient CNS barrier breach, absence of a required ‘second hit’ or low MAIT cell numbers in mice compared to humans (*23, 24*).

In addition to their function in suppressing peripheral autoimmunity, MAIT cells have been shown to accumulate in the inflamed organs in some autoimmune diseases (*25, 26*). We therefore next investigated whether similar mechanisms may apply to AIE. We therefore queried the available rare autopsy materials of 4 VGKC (LGI1-CASPR2-), 4 LGI1+ and 1 CASPR2+ AIE patients for signs of significant T cell infiltration in the parenchyma. Of these cases, two (1 CASPR2 AIE, 1 VGKC AIE) patients showed signs of T cell infiltration which were investigated further by multicolor immunohistochemistry. We identified CD3^+^CD161^+^TCRγδ^-^CD4^-^CD8^-^ and CD3^+^CD161^+^TCRγδ^-^CD4^-^CD8^+^ cells which resemble MAIT cells in both samples in the hippocampus and uncus (Fig. 5F); regions predominantly affected by the disease. Overall, our findings suggest that MAIT cells may exert functions defined by the tissue environment in AIE by peripherally suppressing anti-neuronal autoantibodies and locally promoting tissue damage in the affected CNS as described across diverse other autoimmune conditions (*20, 25–28*)

## Discussion

Here, we combined a deep transcriptomic characterization of blood and CSF cells with histological and functional confirmation to gain novel mechanistic understanding of LGI1- and CASPR2-associated AIE while excluding confounding treatments. We made several main observations: *First*, the single most dominant cellular change in the CSF in LGI1-/CASPR2-AIE was an expansion of plasma cells. Unlike for example multiple sclerosis (MS) that grossly affects diverse CSF lineages (*12*), CSF alterations in LGI1-/CASPR2-AIE were only detected when specifically assessing plasma cells (*29*); i.e. ‘an immunological needle, not a hammer’. *Second*, the plasma cells in CSF were transcribing both IgG2 and IgG4 and clonally expanded with evidence of ongoing affinity maturation within the CSF compartment and almost exclusive autoantigen-specificity. *Third*, T cells in CSF also showed clonal restriction and expansion with some evidence of potentially providing B cell help. *Fourth*, innate-like T cells, especially MAIT cells and γδ T cells were reduced in the CSF and blood of patients with AIE, present in the hippocampi of two autopsy cases and repressed systemic autoantibody production in a murine immunization model. One could thus speculate whether MAIT cells link environmental triggers with anti-neural autoimmunity.

Notably, MAIT cells not only sense microbial patterns in tissue, but also locally shape adaptive antibody responses to infectious pathogens (*20*). Potentially, MAIT cells could lose their autoantibody-suppressive capacity in LGI1-/CASPR2-AIE in direct or indirect contact with danger signals in barrier-tissue, e.g. the gut or possibly the meninges. Indeed a peripheral loss of MAIT cells has been described consistently across autoimmune diseases (*25, 26, 28*) while selected studies show an increase of MAIT cells in the targeted tissue suggesting a compartment-specific function of MAIT cells (*26, 30*). We histologically demonstrate that MAIT cells can indeed home to the site of inflammation in AIE. One could speculate if translocation of innate-like T cells from periphery to target organ could disinhibit peripheral auto-antibody formation. Invariant NK and γδ T cells can provide cognate and noncognate help to B cells in mice (*31–33*) by boosting antibody responses and direct interaction with glycolipids and antigen on CD1d (*34–36*). Furthermore, MAIT cells can promote humoral immunity (*37–40*) by increasing the production and differentiation of plasmablasts in a feed-forward loop (*38, 39*). This link between innate-like T cells and autoantibody-mediated encephalitis variants might identify exploitable therapeutic targets early in the pathogenic cascade of antibody-driven CNS autoimmunity.

We identified several remarkable features of B cell autoimmunity in LGI1-/CASPR2-AIE. *First*, pronounced plasma cell/blast expansion in the CSF was not reflected by more routine surrogates of B lineage pathology in CSF (e.g. OCB, total B cells) as described (*3, 10*). *Second*, expanded clones almost exclusively produced neuronal autoantigen-specific autoantibodies thus arguing against epitope spreading. One study had transcriptionally analyzed 237 single CSF cells from one patient with anti-LGI AIE, but lacked power (*41*). Another previous study comparing LGI1-AIE to NMDAR-AIE did not unequivocally identify autoantigen-specific clones, but also lacked statistical power (*42*). *Third*, expanded clones were almost uniquely present in the CSF with some signs of somatic hypermutation in line with previous results in CSF and blood of six patients with LGI1 encephalitis (*43*). *Fourth*, clones transcribed both IgG2 and IgG4 heavy chain genes, thus arguing against LGI1-/CASPR2-AIE as a strictly IgG4-associated disease. This is in line with a previous study that found 84% of plasma cells producing LGI1-specific antibodies with somatic hypermutation and mainly IgG1,2 and IgG4 heavy chains in three patients undergoing immunotherapy (*44*). *Fifth*, findings in CASPR2-AIE were essentially comparable with LGI1-AIE, thus showing converging mechanisms across AIE subtypes. In accordance, dense perivascular B cell and plasma cell infiltrates have been observed in the meninges and Virchow-Robin spaces (together with CD4 T cells) of patients with LGI1-AIE (*45*) (and NMDAR-AIE (*46*)). Interestingly, the role of meninges in the regulation of CNS immune responses has gained renewed interest (*47*) as an active neuroimmune interface. The meninges might in fact represent an integration point between the adaptive and innate immune system in AIE.

Elucidating this connection between innate-like T cells and B cells necessitated immunization models emulating early components of autoimmunity. Models of passive immunization (*48–51*), including injection with CASPR2 and LGI1 antibodies (*21, 48, 51*), have previously been successfully established in AIE. Naturally, these models only allow studying the effector functions of the respective antibodies. T cell stimulating approaches with vector-based intrathecal application have been published (*52, 53*) for limbic encephalitis, but do not mimic the systemic antigen based response. Recently three similar active immunization methods have been demonstrated in NMDAR-AIE (*22, 54, 55*) that could also show CNS inflammation and behavioral changes mimicking NMDAR-AIE. Thus, murine active immunization models in AIE remain challenging and to the best of our knowledge, we are the first to report any active LGI1/CASPR2 immunization model inducing disease-defining antibodies. Our study may form the basis for mechanistic exploitation of such active immunization models also in LGI1-/CASPR2-related disease and of the role of MAIT-cells in this context

In this study, we provide one of the largest single cell compendiums of CSF cells in any disease. Studies with similar design have accumulated 120,629 (n = 51 donors) (*56*) and 216,723 single cell transcriptomes (175,529 CSF / 41,194 blood; n = 57 donors) from MS patients and controls (*57*). Dementia-related CSF atlassing efforts have collected 22,625 (n = 22 donors) and 21,267 single cell transcriptomes (n = 18 donors) from CSF (*58, 58*). Our present dataset encompasses 162,360 total single-cell transcriptomes from 13 individuals with disease conditions orders of magnitude less prevalent than dementia and multiple sclerosis. The low prevalence of LGI1-/CASPR2-AIE naturally complicated our ability to recruit larger cohorts and to balance confounders between groups. We also provide additional experimental confirmation using additional cohorts and rodent disease modeling whenever possible.

Overall our study provides for the first time a large scale unbiased transcriptomic analysis with confirmation assays in independent cohorts in LGI1 and CASPR2 AIE. In the future it will be crucial to further extend this study to different AIE variants, to longitudinal studies of patients receiving e.g. B cell depleting treatment, and to consider further trans-compartment analyses, e.g. CSF-lymph nodes. Our study also provides potential for mechanistic understanding and diagnostic and therapeutic markers to benefit personalized treatments in autoantibody-mediated brain diseases.

## Methods

### Prospective patient recruitment, CSF collection and retrospective data collection

#### Prospective sampling in cohort 1

All patients with suspected AIE or antibody-confirmed LGI1-/CASPR2-AIE at the centers in Münster, Rotterdam, Kiel were prospectively screened for inclusion into the study. Formal inclusion criteria for AIE patients were: (1) fulfillment of the Gauss criteria for probable AIE, (2) suspected AIE with later antibody confirmed LGI1-/CASPR2-AIE or initially antibody-confirmed LGI1-/CASPR2-AIE, (3) immunotherapy-naive except for a maximum of 1 day of steroid treatment, (4) receiving lumbar puncture (LP) for diagnostic purposes, and (5) written informed consent to participate. All patients were of Caucasian ethnicity. The study was performed in accordance with the declaration of Helsinki and was approved by: (1) the “Ethikkommission der Ärztekammer Westfalen-Lippe (ÄKWL) und der Westfälischen-Wilhelms-Universität” (Ethics Committee of the Board of Physicians of the Region Westfalen-Lippe and of the Westfälische Wilhelms-University Münster) under reference number 2015-522-f-S in Münster, (2) by the “Erasmus MC Medical Ethical Research Board under reference numbers MEC-2015-397, MEC-2020-0418, and MEC-2020-0650 in Rotterdam (3) by the “Ethikkommission” of the medical faculty Kiel reference number: D498/19, B337/13, B300/19, and D578/18.

The IIH patient cohort had partially been reported previously (*13*) or were recruited for the present study and also fulfilled the inclusion criteria 3-5.

Exclusion criteria for all patients in cohort #1 were defined as: (1) questionable diagnosis of AIE by clinical signs or magnetic resonance imaging (MRI) findings, and (2) ongoing or previous immunomodulatory treatment. IIH patients were included, if they gave informed consent. Exclusion criteria for all patients were: (1) immunologically relevant comorbidities (e.g. rheumatologic diseases), (2) severe concomitant infectious diseases (e.g. HIV, meningitis, and encephalitis), (3) pregnancy or breastfeeding, (4) younger than 18 years, (5) mental illness impairing the ability to give informed consent, and (6) artificial blood contamination during the LP resulting in >200 RBCs/μl.

CSF for scRNA-seq was collected as described (*12*). Briefly, during LPs performed for clinical reasons, up to 10 ml of CSF and 3ml of blood were collected in addition to diagnostic material. CSF was transported at 4°C and processed within an hour to ensure optimal sample quality. CSF cells, total protein and intrathecal immunoglobulin concentrations were assessed according to validated standard diagnostic procedures in central labs of each center. Concentration of protein and immunoglobulins in serum and CSF were compared and a Reiber scheme was created to evaluate the integrity of the blood-CSF-barrier (BCBD), quantified by the ratio between CSF albumin and serum albumin. Oligoclonal bands (OCB) were detected by isoelectric focusing and silver nitrate staining. Samples were pseudonymized at collection. CSF was centrifuged at 300×g for 10 min. The supernatant was removed. CSF cells were resuspended in 5ml of X-Vivo15 media (Lonza). A total of 5 μl of the single-cell suspension were manually counted in a counting chamber. The remainder of the CSF cells and a maximum of 20,000 CSF were used as input for scRNA-seq. If total available CSF cell numbers were <17,000 cells, all available cells were processed.

#### Retrospective sampling in cohort 2

We retrospectively screened all patients who were admitted to the University Hospital Münster between 2011 and 2020 and received a diagnostic lumbar puncture, including flow cytometry, for the ICD-10 diagnoses G04.*, G13.1, G93.2. Flow cytometry raw data from all treatment-naive patients was analyzed (Suppl. Fig. 2 A-D).

All CSF cell samples collected during regular working hours at the center in Münster were routinely and promptly analyzed by flow cytometry using a Navios flow cytometer (Beckman Coulter) and an antibody panel described previously (*59*). Briefly, blood cells were lysed using VersaLyse buffer and blood and CSF cells were stained using the following anti-human antibodies (Biolegend; clone names indicated): CD3 (UCHT1); CD4 (13B8.2); CD8 (B9.11); CD14 (RMO52); CD16 (3G8); CD19 (J3-119); CD25 (B1.49.9); CD27 (1A4CD27); CD45 (J.33); CD45RA (ALB11); CD56 (N901, NCAM16.2); CD127 (R34.34); CD138 (B-A38); and HLA-DR (Immu-357). The gating scheme is depicted in (Suppl. Fig. 2A). Cell population size was defined as the number of gated cell events relative to the events of the corresponding parent gate.

#### Retrospective sampling in cohort 3

For blood flow cytometry, we retrospectively screened available cryo-preserved PBMCs of patients with confirmed LGI1-/CASPR2-AIE at the center in Münster. Peripheral blood was cryopreserved, as described previously (*60*). Cryopreserved PBMC were thawed, divided into separate samples and stained with distinct sets of fluorochrome-conjugated antibodies over 30 minutes with anti-human antibodies as indicated. T cell panel consisted of pan-γδTCR (B1), CD3 (OKT3), CD161 (HP3G10), TCRVα7.2 (3C10), CD19(SJ25C1), CD4 (SK2), CD8a (HIT8a), 7-AAD, Tetramer MR1-RU, and MR1-FP(*61*) as control staining. The MR1 tetramer technology was developed jointly by Dr. James McCluskey, Dr. Jamie Rossjohn, and Dr. David Fairlie, and the material was produced by the NIH Tetramer Core Facility as permitted to be distributed by the University of Melbourne. Flow-cytometric data were analyzed using FlowJo V10. Immune cell subsets were defined according to a prespecified gating hierarchy Suppl. Fig. 7B,C,D.

### Single cell RNA-sequencing

Cells were processed for scRNA-seq as previously described (*12, 13*). Briefly, single-cell suspensions were loaded onto the Chromium Single Cell Controller using the Chromium Single Cell GEM, Library & Gel Bead Kits (10X Genomics) with chemistry version V1 and V1.1 for 5’ reagents as well as V3 and V3.1 for 3’ reagents. Sample processing and library preparation was performed according to the manufacturer’s instructions and sequencing carried out on an Illumina Nextseq500 or an Illumina Novaseq 6000 with 2×150 or 2×100 read setup (for details see Suppl. Tab. 2).

### scRNA-seq bioinformatic analyses

The preprocessing of scRNA data was performed with the 10x Genomics’ Cell Ranger software v3.1.0 using the reference GRCh38 v3.0.0. The resulting filtered feature-barcode matrix files were analyzed with the R package Seurat v.4.0.4 and v4.3.0 (*62*). To minimize the number of doublets, empty cells, and cells with a transcriptome in low quality, only cells harboring between 300 and 3000 RNA features and less than 5% mitochondrial RNA were selected for further processing (Suppl. Tab 4). All genes with a detected expression in less than 1% of the cells as well as mitochondrial and ribosomal genes (comprising all gene having names starting with MT-, RPS, RPL or RNA\\d8S5) were not considered for further analyses. Afterwards, data were log-normalized, variable features identified, and features scaled and centered in the dataset. After performing a PCA dimensionality reduction (30 dimensions) with the RunPCA function, the expression values were corrected for effects caused by different library preparation kits (3’ vs 5’) and different sampling locations using the R package Harmony v0.1.1 (*63*). In the final steps, the Uniform Manifold Approximation and Projection (UMAP) dimensional reduction was performed with the RunUMAP function using 15 dimensions, a shared nearest neighbor graph was created with the FindNeighbors function, and clusters were identified with a resolution of 0.2 using the FindClusters function. The clusters were annotated with the R function SingleR (*64*) based on the Human Primary Cell Atlas reference data set followed by a manual correction using marker genes computed with the R function FindMarkers. Two clusters, which were both annotated as monocytes, were merged to one cluster afterwards. To investigate B and T cells, the Seurat object was annotated with the tool Azimuth v0.4.6 using the human PBMC reference (67). Afterwards a separate reclustering was performed for B cells and T cells, respectively. In addition, B-cells were annotated with Azimuth v0.4.6 (59) using the human tonsil (13) reference followed by a manual correction of the cell type assignment.

### Differential gene expression and enrichment analysis

To determine differentially expressed genes (DEG) in a cluster independent fashion, we used the miloDE (*18*) framework v.0.0.0.9000 following the official tutorial. Briefly, we assigned neighborhoods (k = 30, d = 30). Next, DEG was performed within each neighborhood using the de_test_neighbourhoods function with a minimal count of 10. To determine DEG based on the predefined clusters, we used two different approaches: (i) For the figures in the manuscript, differentially expressed genes were determined using the FindMarkers funktion with MAST v1.24.1 (*65*). Thereby, only genes with a minimum fraction of 0.1 in either of the two groups were considered. (ii) For the supplemental tables (T cells and across cluster in CSF), limma v3.54.1 (*66*) was applied. For limma, we generated pseudobulk data removing genes with less than 3 expressing cells and cell types with less than 10 cells per sample. The data were processed with voomWIthQualityWeights and a linear model was fit, followed by an empirical Bayes smoothing to determine DE genes. The gene set enrichment analyses were performed with the online tool InnateDB (*67*) based on entries from the KEGG (*68*) database. For visualization, obvious disease-associated pathways were excluded to remove artifacts. Ligand-receptor interactions between B and T cells were calculated with the R tool CellChat v1.6.1 based on the subset ‘Secreted Signaling’ of the human CellChatDB database and the ‘weight’ option (*70*). For this analysis, mature plasma cell types as well as plasma cell precursor cell types were merged, respectively.

### Cell abundance analysis

We used the propeller method (*71*) (part of speckle v0.99.1) to determine differentially abundant clusters across all cell types and in the T cell analyses. We removed clusters with less than 30 cells in both conditions combined in advance. Following the official tutorial, a logit transformation was performed. Next, a t-test was performed with the propeller.ttest function with a robust variance estimation.

### BCR and TCR bioinformatic analysis

The preprocessing of the VDJ sequencing data was performed with 10x Genomics’ Cell Ranger software v3.1.0 with the VDJ reference GRCh38 v3.1.0. Based on the Cell Ranger output, the Gini-Simpson diversity was determined with the diversity function of the R microbiome package v1.23.1, while the shannon and the inverse simpson index were calculated with the diversity function of the R vegan package v2.6.4. The reported cell frequencies for the clones were visualized in the UMAP plots. In the BCR networks, cells were connected, if they had the same variable, joining, and constant gene as well as the same CDR3 region on the IGH chain. In the TCR networks, cells were connected, if they had the same variable, diversity, joining, and constant gene segments as well as the same CDR3 region on the TRB chain. To determine the somatic hypermutation, clones with at least three cells were extracted and the consensus sequence of each clone aligned with all human IGHV reference sequences annotated in the IMGT database (*72*). The reference sequence with the highest identity was then compared with all sequences of the cells and the mean number of mutations per clone determined. In addition, we performed for each clone a hierachical clustering using the R function hclust out of the stats package v4.2.2. TCR sequence sharing was assessed with GLIPH v2 (*73, 74*) and TCR antigen association with VDJdb (*75*).

### Mouse immunisation

All animal experiments were approved by the local authorities (Landesamt für Natur, Umwelt und Verbraucherschutz Nordrhein-Westfalen; Approval ID: 84-02.04.2022.A336). Every effort was made to minimize the number of animals used and to avoid stress and suffering of the animals by strictly following the ARRIVE guidelines. The mice were housed in groups, had a 12-hour light/dark cycle, and food and water were available ad libitum. Homozygous MR1 deficient (MR1AIE) and C57BL/6 (C57BL6AIE) mice (10-15 weeks old, male and female) were immunized twice with Recombinant Mouse CNTNAP (>95% purity) and LGI1 protein (>90% purity) (Cntnap2-3316M, LGI1-9069M, Creative Biomart) emulsified in Complete Freund’s Adjuvant (CFA) and supplemented with Mycobacterium tuberculosis H37Ra (4 mg/mL). Mice were immunized subcutaneously on the back with 100 μg of the protein in the emulsion mixture. Mice in the control groups, including homozygous MR1 deficient (MR1control) and C57BL/6 (C57BL6control) mice, received an emulsion mixture of CFA and the same volume of phosphate-buffered saline (PBS). All mice were injected intraperitoneally with 250 ng of pertussis toxin (Sigma) on the day of immunization and 48 hours later. Serum from all mice was collected on the 15th and 28th day to determine antibody titers.

### Murine evaluation

Mice were evaluated 3-1 days prior, at 12-14 days and at 25-28 days after immunization and underwent behavioral tests to evaluate locomotor activity, anxiety levels and memory. All tests were evaluated by an automated program (Noldus Ethovision (*76*)) with manual correction. The Open Field test assessed locomotor activity and exploratory behavior of mice (*77*) Animals were tested in the open field arena and traveled distance, velocity, and time spent in the center were measured.

The Novel object recognition test (NOR) (*78, 79*) was used to assess cognitive and memory functions. The same arena was used as animals were already familiar with it. After 3 familiarization sessions over 3 days animals were able to explore the arena with 2 identical objects for 5 minutes. Two test phases were tested to evaluate short term (1 hour) and long term (24 hours) memory (*80*). For the testing session one of the familiar objects was replaced by a novel object. The time spent exploring the novel and old object was used to calculate the Novel object recognition index (NOR Index: (time novel/ (time novel+time old)).

The three-chamber test (*81, 82*) was performed in a plastic apparatus (60×60cm) with 3 chambers delimited by removable dividers after habituation a day prior. Testing was performed in 2 phases. Phase 1 included a 10 minute acclimation phase with 2 empty cages in the outer chambers. In phase 2 an empty cage(non-social stimulus) and a cage with an unfamiliar mouse (of the same sex) (social stimulus) was placed in the outer chambers.Time spent in each chamber was recorded over 10 minutes. The social preference index was calculated (social stimulus-non social stimulus/(social stimulus+nonsocial stimulus).

Mouse Serum of day 14 and day 28 was extracted from the facial vein, centrifuged for 10 minutes and analyzed for the presence of IgG autoantibodies against neural surface membrane antigens (NMDAR, AMPAR, GABA, LGI1, CASPR2, DPPX) using a cell-based assay according to manufacturer’s instructions (EUROIMMUN, Lübeck, Germany) with a secondary Anti-mouse-IgG Antibody (Biolegend, Poly4053). Dilution steps included pure, 1:1, 1:10, 1:32, 1:100,1:1000 according to standard clinical testing. Positivity for fluorescence was determined blinded on a Zeiss Axioscope (AX10).

Mice were perfused intracardially. The brain was extracted, digested with collagenase D (2.5 mg/mL) and DNase I (0.05 mg/mL) (20 min, 37 °C) and leukocytes were purified using a 70/37% Percoll gradient. Cells were stained for flow cytometry with anti-mouse antibodies (Biolegend; clone names indicated): CD45(30-F11), CD3 (17A2), CD4 (GK1.5), gdTCR (GL3), NK1.1 (S17016D) and CD11b (M1/70). Live/dead staining was performed with Zombie NIR. Anti-Mouse MR1 Tetramer MR1-RU (*61*) and control MR1-FP stainings were performed. Tetramer gating was introduced through a negative control for each sample (Suppl. Fig. 8C).The MR1 tetramer technology was developed jointly by Dr. James McCluskey, Dr. Jamie Rossjohn, and Dr. David Fairlie, and the material was produced by the NIH Tetramer Core Facility as permitted to be distributed by the University of Melbourne. Cells were analyzed using a Aria flow cytometer (Beckman Coulter). Data were analyzed with FlowJo v10

### Flow cytometric and murine statistics

Significance was tested with a Kruskal Wallis and post hoc Dunn test to compare multiple groups and the Benjamini Hochberg method used for multiple testing corrections. If two groups were compared a Mann-Whitney U test was used. Overall Fishers test was used to show significance in antibody incidence and post hoc individual Fishers test were performed to assess subgroup significance. The computational analysis was carried out with R 4.1.1.

### Human CNS multiplex immunofluorescence labeling

Histological paraffin sections of brain tissue from autopsied patients (n=2) with the diagnosis CASPR2-AIE and anti-VGKC AIE were included. Sections were localized in the uncal and hippocampal region. Immunofluorescence labeling of sections was performed using markers for CD3(Neomarkers, #RM9107-S), CD8 (Dako M7103, CD4 (Cell Signailing, #48274), TCR δ(SantaCruz, #sc-100289) and CD161 (Abcam, #ab302564). The staining procedure was executed in accordance with the Akoya Fluorescent Multiplex kit protocol (*83, 84*). In short, antigen retrieval was achieved by placing samples in EDTA pH9 for 60 minutes in a household food steamer (Braun), followed by a 10-minute incubation with Opal Antibody Dilutent/Block solution (Akoya Bioscience, Marlborough, USA). The first primary antibody was then applied overnight at 4°C. Subsequently, sections were rinsed several times in Tris-buffer saline with Tween 20 (TBS-T) and the secondary antibody was applied. Hence either horseradish peroxidase (HRP) conjugated donkey α-mouse (Jackson, #715-225-151) or HRP conjugated donkey α-rabbit (Jackson, #711-035-152) was used. Afterwards, one of the fluorophores was introduced (Opal 570, Opal 690, Opal 480, Opal 620). Before proceeding with the next primary antibody, the samples were fixed with 4% paraformaldehyde for 10 minutes at RT followed by another antigen retrieval step with AR6 for 30 minutes. Regarding Opal 780, the staining procedure differed: after incubation with the secondary antibody, Opal TSA-DIG (Akoya Bioscience, Marlborough, USA) was introduced for 10 minutes. Next, the sections were transferred into AR6 and heated in a household food steamer for 20 minutes. The fluorophore signal was generated by incubating the sections with Opal 780 at RT for 60 minutes and 4′,6-diamidino-2-phenylindole (DAPI) was applied for counterstaining.

To quantify, cells were scanned with the Vectra Polaris Automated Quantitative Pathology Imaging system from Perkin Elmer and quantified with Qupath software. To this end, cells were detected by nuclear staining (DAPI).

### Cloning antibodies and verification of antigen-specificity

Corresponding full length consensus sequences of variable heavy (VH) and light chains (VL) including signal peptide sequences of the top 5-9 most abundant, clonally expanded B cells in the CSF sample of all patients were determined from 5’ VDJ 10x single cell information. Cloning strategy, vectors, and expression/purification strategies were kindly provided by M. Peipp, Kiel and done as previously described (*85*). In brief, for VH sequences, N-terminal NheI and C-terminal PpuMI sites (gcctccactaagggacct) and for kappa and lambda chains, NheI und NotI sites were added. All sequences were synthesized by GeneArt (ThermoFisher) and cloned into pcDNA3.1(+) expression vectors. Heavy chain variable regions were subcloned using NheI/PpuMI into an acceptor vector containing a IgG4 heavy constant region. Human Embryonic Kidney (HEK-293) cells were transfected with Lipofectamine 2000 (Invitrogen 11668-019) and 16 μg DNA per vector (light and heavy chains). Supernatant was incubated with CaptureSelect™ IgG-CH1 Affinity Matrix (Thermo Fisher 1943200250) and antibodies were eluted by acidic elution with (pH 2.8) and neutralized. Purified recombinant human antibodies (rHumAbs) were desalted (Zeba™ Spin Desalting Columns, Thermo Fisher 89891) and concentrated using a Christ Rotational Vacuum Concentrator System. Antibody concentration was determined with Qbit protein assay (Invitrogen Q33211).

Antigen-specificity was confirmed using cell-based assays (CBA). HEK-293 cells were transfected with a plasmid encoding full-length human LGI1 c-terminally fused to a transmembrane CASPR2 domain and intracellular GFP, or full-length human CASPR2 with C-terminal GFP tag (both kindly provided by S. Irani, Oxford), using Lipofectamine 2000. After overnight incubation at 4 °C, cells were incubated with rHumAbs (0.4 μg/mL) at 37 °C for 60 minutes, fixed with 4% PFA at 4 °C for 5 minutes, permeabilized with 0.3% Triton X-100 (Th-Geyer 8059.0250) for 5 minutes, and blocked with 1% bovine serum albumin (BSA) for 90 minutes. Secondary staining was done using a goat anti-Human Alexa Fluor (AF) 594 antibody (1:1000, Invitrogen), for 30 minutes. Cells were mounted in Vectashield HardSet Mounting Medium with DAPI (4′,6-diamidino-2-phenylindole, Vector H-1500), and results were evaluated using a fluorescence microscope. Human serum (1:40) from autoimmune encephalitis patients or healthy subjects was used as positive and negative controls, respectively.

## Acknowledgments

We thank all participating patients and their relatives. We thank the GENERATE network as a general framework of researchers interested in autoimmune encephalitis in Germany. FL and MT are members of the European Reference Network ERN-RITA.

## Author contributions

DE, LMM, MH, MP, MVD, LAC, KM, SR, EB, and JD performed data acquisition. DE, LMM, and MH carried out data analysis. FL and GMzH conceived and supervised the study. HW, SGM, JB, MT, and NM co-supervised the study. DE, LMM, FL, and GMzH wrote the manuscript. All authors read and approved the final version of the manuscript.

## Data and materials availability

The sequencing raw data and processed data will be made available in the NIMH data archive upon reasonable request and after material transfer agreement to regulate data protection of potentially re-identifiable genetic data.

